# Evolution of microbial growth traits under serial dilution

**DOI:** 10.1101/798678

**Authors:** Jie Lin, Michael Manhart, Ariel Amir

**Affiliations:** John A. Paulson School of Engineering and Applied Sciences, Harvard University, Cambridge, MA 02138, USA; Department of Chemistry and Chemical Biology, Harvard University, Cambridge, MA 02138, USA; Institute of Integrative Biology, ETH Zurich, 8092 Zurich, Switzerland

## Abstract

Selection of mutants in a microbial population depends on multiple cellular traits. In serial-dilution evolution experiments, three key traits are the lag time when transitioning from starvation to growth, the exponential growth rate, and the yield (number of cells per unit resource). Here we investigate how these traits evolve in laboratory evolution experiments using a minimal model of population dynamics, where the only interaction between cells is competition for a single limiting resource. We find that the fixation probability of a beneficial mutation depends on a linear combination of its growth rate and lag time relative to its immediate ancestor, even under clonal interference. The relative selective pressure on growth rate and lag time is set by the dilution factor; a larger dilution factor favors the adaptation of growth rate over the adaptation of lag time. The model shows that yield, however, is under no direct selection. We also show how the adaptation speeds of growth and lag depend on experimental parameters and the underlying supply of mutations. Finally, we investigate the evolution of covariation between these traits across populations, which reveals that the population growth rate and lag time can evolve a nonzero correlation even if mutations have uncorrelated effects on the two traits. Altogether these results provide useful guidance to future experiments on microbial evolution.

---

Laboratory evolution experiments in microbes have provided insight into many aspects of evolution [1–3], such as the speed of adaptation [4], nature of epistasis [5], the distribution of selection coefficients from spontaneous mutations [6], mutation rates [7], the spectrum of adaptive genomic variants [8], and the preponderance of clonal interference [9]. Despite this progress, links between the selection of mutations and their effects on specific cellular traits have remained poorly characterized. Growth traits — such as the lag time when transitioning from starvation to growth, the exponential growth rate, and the yield (resource efficiency) — are ideal candidates for investigating this question. Their association with growth means they have relatively direct connections to selection and population dynamics. Furthermore, highthroughput techniques can measure these traits for hundreds of genotypes and environments [10–13]. Numerous experiments have shown that single mutations can be pleiotropic, affecting multiple growth traits simultaneously [14, 15]. More recent experiments have even measured these traits at the single-cell level, revealing substantial non-genetic heterogeneity [10, 13, 16]. Several evolution experiments have found widespread evidence of adaptation in these traits [17–20]. This data altogether indicates that covariation in these traits is pervasive in microbial populations.

There have been a few previous attempts to develop quantitative models to describe evolution of these traits. For example, Vasi et al. [17] considered data after 2000 generations of evolution in *Escherichia coli* to estimate how much adaptation was attributable to different growth traits. Smith [21] developed a mathematical model to study how different traits would allow strains to either fix, go extinct, or coexist; Wahl and Zhu [22] focused on how the fate of different trait-affecting mutations was determined by their time of occurrence during the growth cycle. However, simple quantitative results that can be used to interpret experimental data have remained lacking. More recent work [23, 24] derived a quantitative relation between growth traits and selection, showing that selection consists of additive components on the lag and growth phases. However, this did not address the consequences of this selection for evolution, especially the adaptation of trait covariation.

In this work we investigate a model of evolutionary dynamics with covariation across multiple growth traits. We consider a minimal model in which different strains of cells interact only by competition for a single limiting resource. We find that the fixation probability of a mutation, even in the presence of substantial clonal interference, is accurately determined by a linear combination of its change in growth rate and change in lag time relative to its immediate ancestor; the relative weight of these two components is determined by the dilution factor. Yield, on the other hand, is under no direct selection. We provide quantitative predictions for the speed of adaptation of growth rate and lag time as well as their evolved covariation. Specifically, we find that even in the absence of an intrinsic correlation between growth and lag due to mutations, these traits can evolve a nonzero correlation due to selection and variation in number of fixed mutations.

## METHODS

### Model of population dynamics

We consider a model of asexual microbial cells in a well-mixed batch culture, where the only interaction between different strains is competition for a single limiting resource [23, 24]. Each strain *i* is characterized by a lag time *L*_*i*_, growth rate *r*_*i*_, and yield *Y*_*i*_ (see Fig. 1a for a two-strain example). Here the yield is the number of cells per unit resource [17]. Note that some of our notation differs from related models in previous work, some of which used *g* for growth rate and *λ* for lag time [23], while others used *λ* for growth rate [25]. Although it is possible to extend the model to account for additional growth traits such as a death rate or lag and growth on secondary resources, here we focus on the minimal set of traits most often measured in microbial phenotyping experiments [10–12, 14–16, 18, 26].

**FIG. 1.**
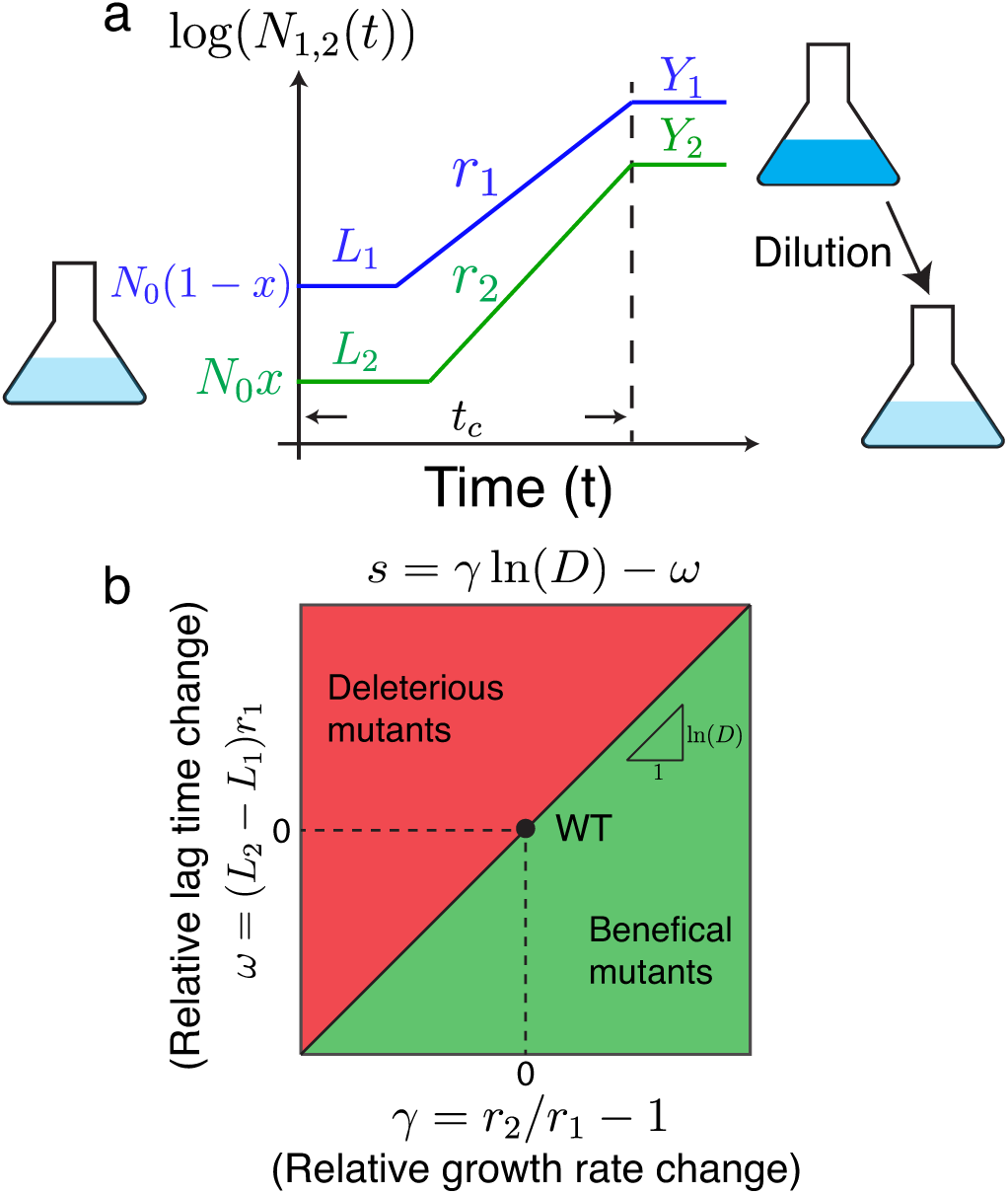
Model of selection on multiple microbial growth traits. (a) Simplified model of microbial population growth characterized by three traits: lag time *L*, growth rate *r*, and yield *Y*. The total initial population size is *N*_0_ and the initial frequency of the mutant (strain 2) is *x*. After the whole population reaches stationary phase (time *t*_c_), the population is diluted by a factor *D* into fresh media, and the cycle starts again. (b) Phase diagram of selection on mutants in the space of their growth rate *γ* = *r*_2_*/r*_1_ *−* 1 and lag time *ω* = (*L*_2_ *− L*_1_)*r*_1_ relative to a wild-type. The slope of the diagonal line is ln *D*.

When the population has consumed all of the initial resource, the population reaches stationary phase with constant size. The time *t*_c_ at which this occurs is determined by equating the total amount of resources consumed by the population at that time with the total initial amount of resources *R*:

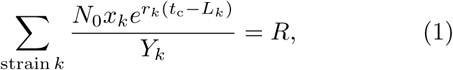

where *N*_0_ is the total population size and *x*_*k*_ is the frequency of each strain *k* at the beginning of the growth cycle. In Eq. 1 we assume the time *t*_c_ is longer than each strain’s lag time *L*_*k*_. We define the selection coefficient between each pair of strains as the change in their log-ratio over the complete growth cycle [27, 28]:

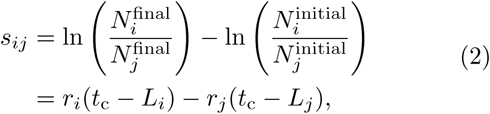

where 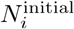 is the population size of strain *i* at the beginning of the growth cycle and 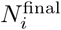 is the population size of strain *i* at the end. After the population reaches stationary phase, it is diluted by a factor of *D* into a fresh medium with amount *R* of the resource, and the cycle repeats (Fig. 1a). We assume the population remains in the stationary phase for a sufficiently short time such that we can ignore death and other dynamics during this phase [29, 30].

Over many cycles of growth, as would occur in a laboratory evolution experiment [1, 28, 31], the population dynamics of this system are characterized by the set of frequencies {*x*_*k*_} for all strains as well as the matrix of selection coefficients *s*_*ij*_ and the total population size *N*_0_ at the beginning of each cycle. In Supplementary Methods we derive explicit equations for the deterministic dynamics of these quantities over multiple cycles of growth for an arbitrary number of strains. In the case of two strains, such as a mutant and a wild-type, the selection coefficient is approximately

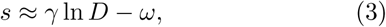

where *γ* = (*r*_2_ *− r*_1_)*/r*_1_ is the growth rate of the mutant relative to the wild-type and *ω* = (*L*_2_ *− L*_1_)*r*_1_ is the relative lag time. The approximation is valid as long as the growth rate difference between the mutant and the wile-type is small, which is true for most single mutations [6, 32]. This equation shows that the growth phase and lag phase make distinct additive contributions to the total selection coefficient, with the dilution factor *D* controlling their relative magnitudes (Fig. 1b). This is because a larger dilution factor will increase the amount of time the population grows exponentially, hence increasing selection on growth rate. Neutral coexistence between multiple strains is therefore possible if these two selection components balance (*s* = 0), although it requires an exact tuning of the growth traits with the dilution factor (Supplementary Methods) [23, 24]. With a fixed dilution factor *D*, the population size *N*_0_ at the beginning of each growth cycle changes according to (Supplementary Methods):

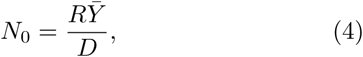

where 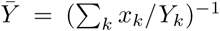 is the effective yield of the whole population in the current growth cycle. In this manner the ratio *R/D* sets the bottleneck size of the population, which for serial dilution is approximately the effective population size [31], and therefore determines the strength of genetic drift.

### Model of evolutionary dynamics

We now consider the evolution of a population as new mutations arise that alter growth traits. We start with a wild-type population having lag time *L*_0_ = 100 and growth rate *r*_0_ = (ln 2)*/*60 ≈ 0.012, which are roughly consistent with *E. coli* parameters where time is measured in minutes [17, 31]; we set the wild-type yield to be *Y*_0_ = 1 without loss of generality. As in experiments, we vary the dilution factor *D* and the amount of resources *R*, which control the relative selection on growth versus lag (set by *D*, Eq. 3) and the effective population size (set by *R/D*, Eq. 4). We also set the initial population size to its steady state value of *N*_0_ = *RY*_0_*/D* (Supplementary Methods).

The population grows according to the dynamics in Fig. 1a. Each cell division can generate a new mutation with probability *µ*, which we set to *µ* = 10^*−*6^; note this rate is only for mutations altering growth traits, and therefore it is lower than the rate of mutations anywhere in the genome. We therefore generate a random waiting time *τ*_*k*_ for each strain *k* until the next mutation with instantaneous rate *µr*_*k*_*N*_*k*_(*t*). When a mutation occurs, the growth traits for the mutant are drawn from a distribution *p*_mut_(*r*_2_, *L*_2_, *Y*_2_*|r*_1_, *L*_1_, *Y*_1_), where *r*_1_, *L*_1_, *Y*_1_ are the growth traits for the background strain on which the new mutation occurs and *r*_2_, *L*_2_, *Y*_2_ are the traits for the new mutant. We will assume mutational effects are not epistatic and scale with the trait values of the background strain, so that *p*_mut_(*r*_2_, *L*_2_, *Y*_2_*|r*_1_, *L*_1_, *Y*_1_) = *p*_mut_(*γ, ω, δ*), where *γ* = (*r*_2_ *− r*_1_)*/r*_1_, *ω* = (*L*_2_ *− L*_1_)*r*_1_, and *δ* = (*Y*_2_ *− Y*_1_)*/Y*_1_ (Supplementary Methods). For simplicity, we focus on uniform distributions of mutational effects where *−* 0.02 *< γ <* 0.02, *−* 0.05 *< ω <* 0.05, and *−* 0.02 *< δ <* 0.02, but in Supplementary Methods we extend our main results to the case of Gaussian distributions as well. Note that since mutations only arise during the exponential growth phase, beneficial or deleterious effects on lag time are not realized until the next growth cycle [20]. After the growth cycle ceases (once the resource is exhausted according to Eq. 1), we randomly choose cells, each with probability 1*/D*, to form the population for the next growth cycle.

## RESULTS

### Fixation of mutations

We first consider the fixation statistics of new mutations in our model. In Fig. 2a we show the relative growth rates *γ* and relative lag times *ω* of fixed mutations, along with contours of constant selection coefficient *s* from Eq. 3. As expected, fixed mutations all increase growth rate (*γ >* 0), decrease lag time (*ω <* 0), or both. In contrast, the yield of fixed mutations is the same as the ancestor on average (Fig. 2d); indeed, the selection coefficient in Eq. 3 does not depend on the yields (Supplementary Methods). If a mutation arises with significantly higher or lower yield than the rest of the population, the bottleneck population size *N*_0_ immediately adjusts to keep the overall fold-change of the population during the growth cycle fixed to the dilution factor *D*. Therefore mutations that significantly change yield have no effect on the overall population dynamics.

**FIG. 2.**
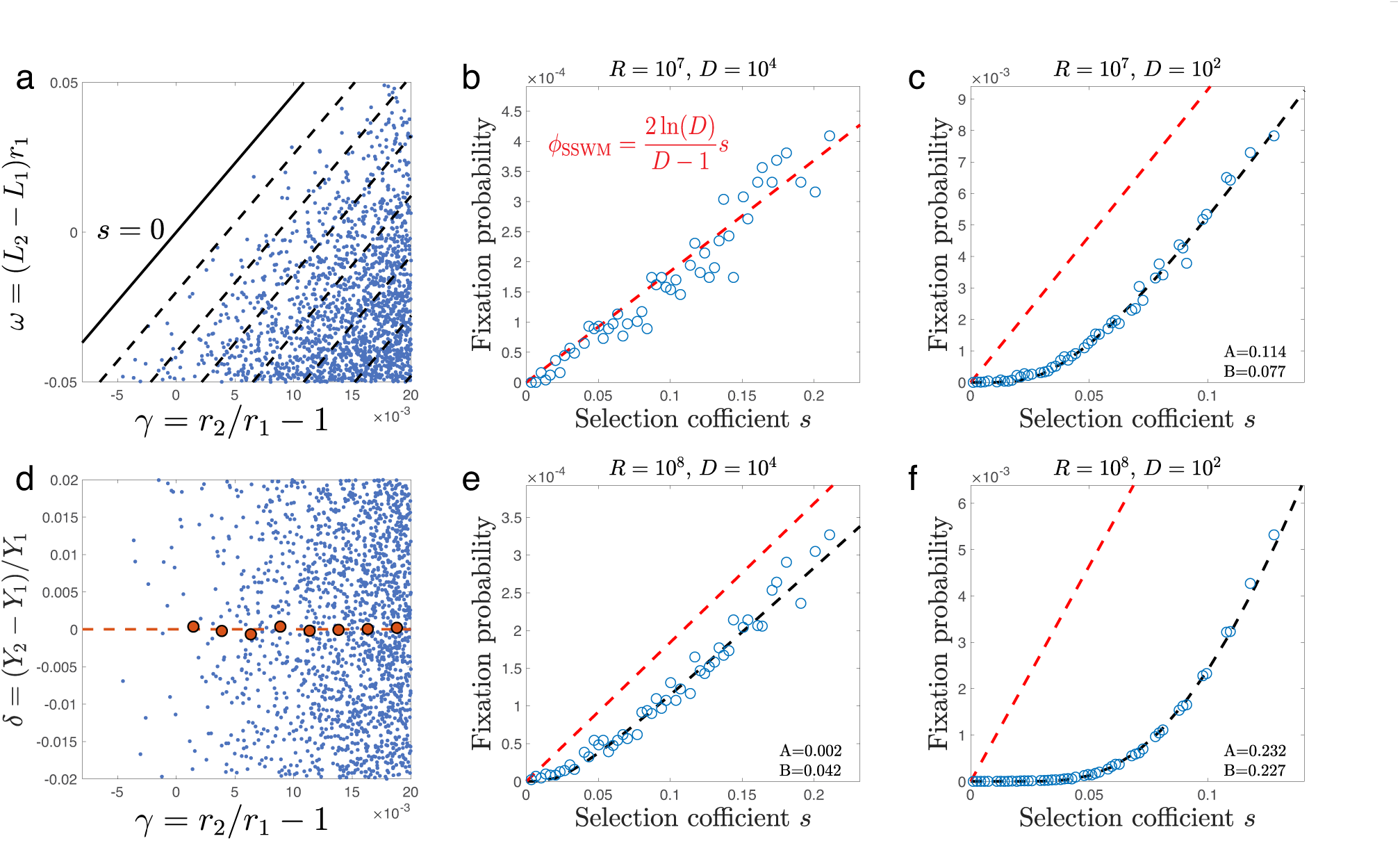
Selection coefficient determines fixation probability. (a) The relative growth rate *γ* and lag time *ω* of fixed mutations. Dashed lines mark contours of constant selection coefficient, while the solid line marks *s* = 0. (d) Same as (a) but for relative growth rate *γ* and relative yield *δ*. The red dots mark the relative yield of fixed mutations averaged over binned values of the relative growth rate *γ*. In (a) and (d), *D* = 10^2^ and *R* = 10^7^. (b,c,e,f) Fixation probability of mutations against their selection coefficient for different amounts of resource *R* and dilution factors *D* as indicated in the titles. The red dashed line shows the fixation probability predicted in the SSWM regime (Eq. 5), while the black line shows a numerical fit of the data points to the fixation probability under clonal interference (Eq. 6), with the resulting fitting parameters *A* and *B* shown in the lower right corner of each panel. In all panels mutations randomly arise from a uniform distribution *p*_mut_ (Supplementary Methods).

Figure 2a also suggests that the density of fixed mutations depends only on their selection coefficient *s*, rather than their individual combination of traits. We therefore plot the fixation probabilities of mutations as functions of their selection coefficients calculated by Eq. 3 (Fig. 2 b,c,e,f). We test this over a range of population dynamics regimes by varying the dilution factor *D* and the amount of resources *R*. For small populations, mutations generally arise and either fix or go extinct one at a time, a regime known as “strong-selection weak-mutation” (SSWM) [33]. In this case, we expect the fixation probability of a beneficial mutation with selection coefficient *s >* 0 to be [22, 34, 35]

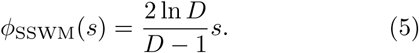

This is similar to the standard Wright-Fisher fixation probability of 2*s* [36], but with a correction to account for the fact that the mutation can arise at different times in the exponential growth phase. Indeed, we see this predicted dependence matches the simulation results for the small population size of *N*_0_ *∼ R/D* = 10^3^ (Fig. 2b).

For larger populations, multiple beneficial mutations will be present simultaneously in the population and interfere with each other, an effect known as clonal interference [37, 38]. We find that the fixation probability of a mutant in this clonal interference regime is well fit by

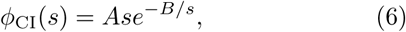

where *A* and *B* are two constants that depend on other parameters of the population; we treat these as empirical parameters to fit to the simulation results, although Gerrish and Lenski [37] predicted *A* = 2 ln *D/*(*D −* 1), i.e., the same constant as in the SSWM case (Eq. 5). Conceptually, this means that interference from other beneficial mutations suppresses the SSWM fixation probability by an exponential factor, where the 1*/s* term comes from the time between the appearance of mutation and its fixation [37]. Equation 6 matches our simulation results well for larger population sizes *N*_0_ *∼ R/D >* 10^4^ (Fig. 2c,e,f). Furthermore, the constant *A* we fit to the simulation data is indeed close to the predicted value of 2 ln *D/*(*D −* 1), except in the most extreme case of *N*_0_ *∼ R/D* = 10^6^ (Fig. 2f).

Altogether Fig. 2 shows that mutations with different effects on cell growth — for example, a mutant that increases growth rate and a mutant that decreases lag time — can nevertheless have the same fixation probability as long as their overall effect on selection is the same according to Eq. 3. In Supplementary Methods we show that these results also hold for a Gaussian distribution of mutational effects *p*_mut_(*γ, ω, δ*), including the presence of correlated mutational effects (Fig. S1). While the dependence of fixation probability on the selection coefficient is a classic result of population genetics [39], the existence of a simple relationship here is nontrivial since, strictly speaking, selection in this model is not only frequency-dependent [23] (i.e., selection between two strains depends on their frequencies) but also includes higher-order effects [24] (i.e., selection between strain 1 and strain 2 is affected by the presence of strain 3). Therefore in principle, the fixation probability of a mutant may depend on the specific state of the population in which it is present, while the selection coefficient in Eq. 3 only describes selection on the mutant in competition with its immediate ancestor. However, we see that, at least for the parameters considered in our simulations, these effects are negligible in determining the eventual fate of a mutation.

### Adaptation of growth traits

As Fig. 3a shows, many mutations arise and fix over the timescale of our simulations, which lead to predictable trends in the quantitative traits of the population. We first determine the relative fitness of the population against the ancestral strain by simulating competition between an equal number of evolved and ancestral cells for one cycle, analogous to common experimental measurements [1, 31]. The resulting fitness trajectories are shown in Fig. 3b. To see how different traits contribute to the fitness increase, we also calculate the average population traits at the beginning of each cycle, e.g., *r*_pop_(*n*) = ∑*i r*_*i*_*/N*_0_(*n*) (where the sum is over all cells), as a function of the number *n* of growth cycles. As expected from Eq. 3, the average growth rate increases (Fig. 3c) and the average lag time decreases (Fig. 3d) for all simulations. In contrast, the average yield evolves without apparent trend (Fig. 3e), since Eq. 3 indicates no direct selection on yield.

**FIG. 3.**
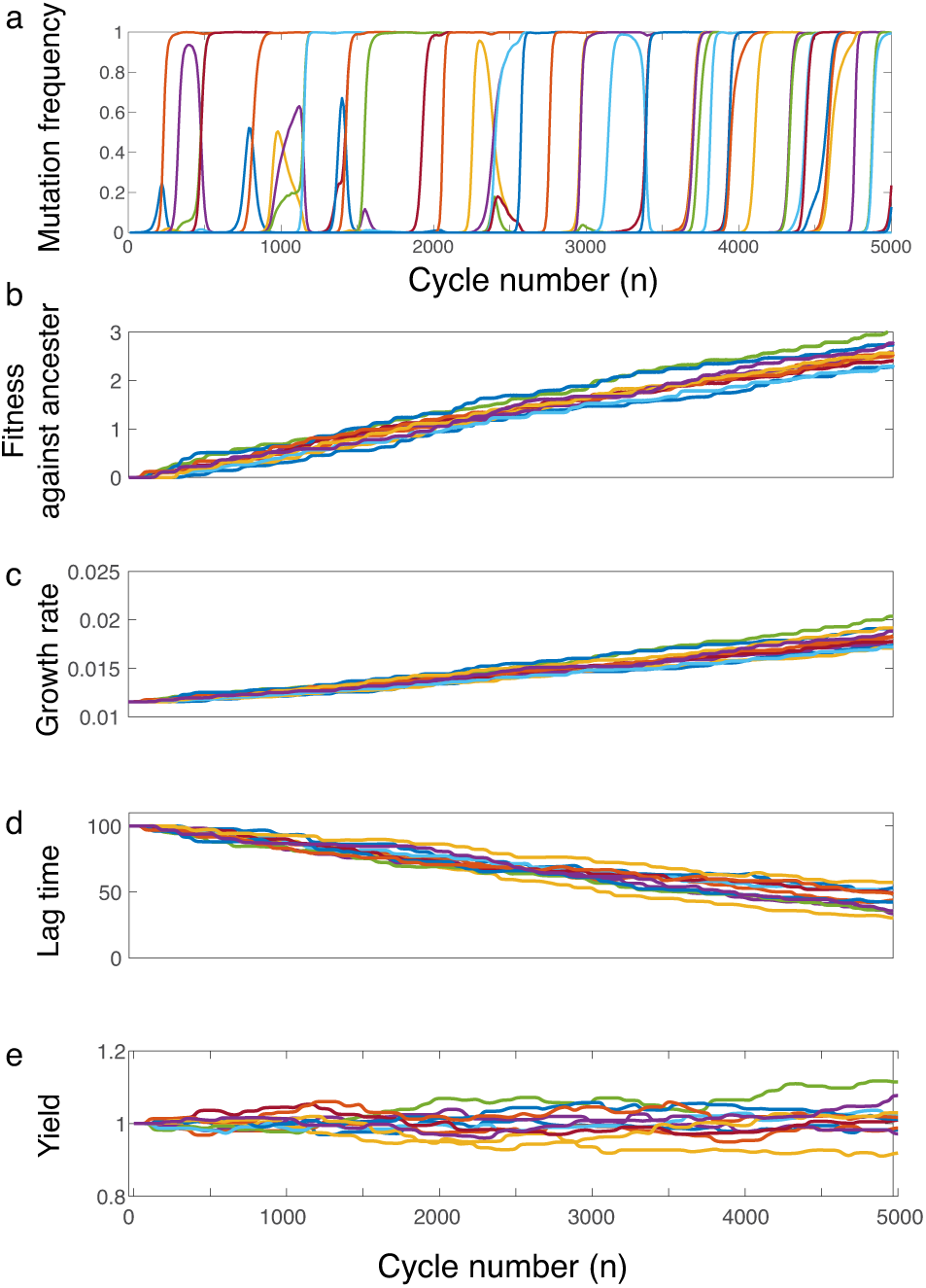
Dynamics of evolving populations. (a) Frequencies of new mutations as a function of the number *n* of growth cycles. Example trajectories of (b) the fitness of the evolved population relative to the ancestral population, (c) the evolved population growth rate, (d) the evolved population lag time, and (e) the evolved population yield. In all panels the dilution factor is *D* = 10^2^, the amount of resource at the beginning of each cycle is *R* = 10^7^, and mutations randomly arise from a uniform distribution *p*_mut_ (Supplementary Methods).

Figure 3 suggests relatively constant speeds of adaptation for relative fitness, growth rate, and lag time. For example, we can calculate the adaptation speed of growth rate as the average change in population growth rate per cycle:

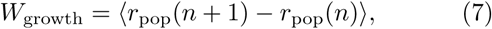

where the bracket denotes an average over replicate populations and cycle number. In Supplementary Methods we calculate the adaptation speeds of these traits in the SSWM regime to be

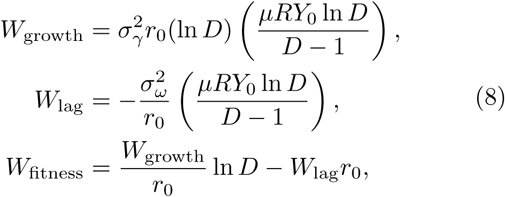

where *σ*_*γ*_ and *σ*_*ω*_ are the standard deviations of the underlying distributions of *γ* and *ω* for single mutations (*p*_mut_(*γ, ω, δ*)), and *r*_0_ is the ancestral growth rate and *Y*_0_ the ancestral yield (which we assume does not change on average according to Fig. 3e). Note the adaptation speeds are proportional to the variances of the traits, which is formally similar to the multivariate breeder’s equation [40], Fisher’s fundamental theorem of natural selection [41], and the Price equation [42]; however, in our case these are variances across single mutants in the SSWM regime, rather than variances of traits within a population. Furthermore, the ratio of growth and lag adaptation rates is independent of the amount of resource and mutation rate in the SSWM regime:

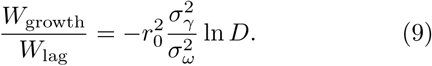

Equation 8 predicts that the adaptation speeds of growth rate, lag time, and relative fitness should all increase with the amount of resources *R* and decrease with the dilution factor *D* (if *D* is large); even though this prediction assumes the SSWM regime (relatively small *N*_0_ *∼ R/D*), it nevertheless holds across a wide range of *R* and *D* values (Fig. 4a,b,c), except for *R* = 10^8^ where the speed of fitness increase is non-monotonic with *D* (Fig. 4c). The predicted adaptation speeds in Eq. 8 also quantitatively match the simulated trajectories in the SSWM case (Fig. 4d,e,f); even outside of the SSWM regime, the relative rate in Eq. 9 remains a good prediction at early times (Fig. S2).

**FIG. 4.**
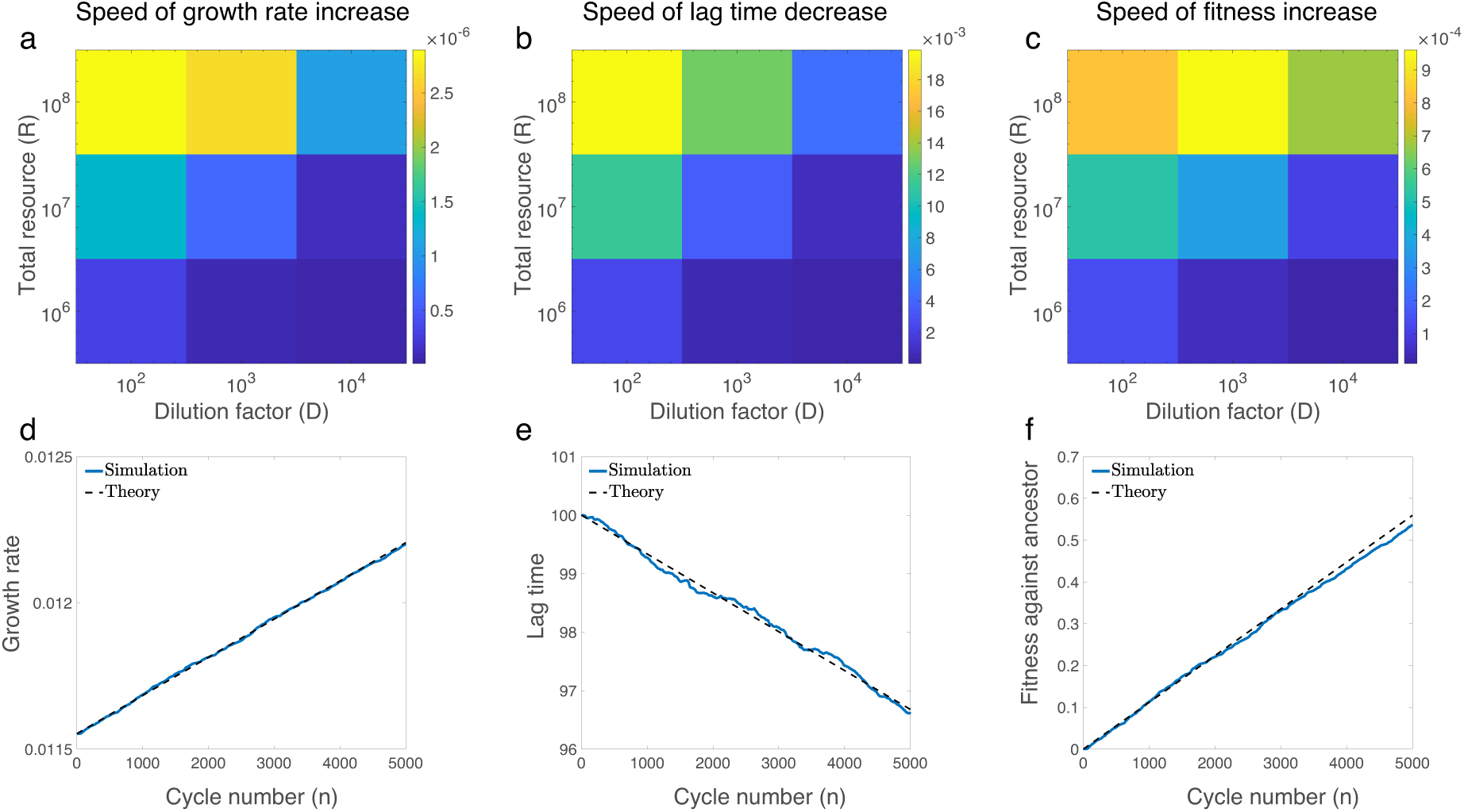
Speed of adaptation. The average per-cycle adaptation speed of (a) growth rate, (b) lag time, and (c) fitness relative to the ancestral population as functions of the dilution factor *D* and total amount of resources *R*. The adaptation speeds are averaged over growth cycles and independent populations. Average population values of (d) growth rate, (e) lag time, and (f) fitness relative to the ancestral population as functions of the number *n* of growth cycles. The dilution factor is *D* = 10^4^ and the total resource is *R* = 10^7^, so the population is in the SSWM regime. The blue solid lines are simulation results, while the dashed lines show the mathematical predictions in Eq. 8. All panels show averages over 500 independent simulated populations, with mutations randomly arising from a uniform distribution *p*_mut_ in which *−*0.02 *< γ <* 0.02, *−*0.05 *< ω <* 0.05, *−*0.02 *< δ <* 0.02 (Supplementary Methods).

### Evolved covariation between growth traits

We now turn to investigating how the covariation between traits evolves. We have generally assumed that individual mutations have uncorrelated effects on differentraits (Supplementary Methods). Nevertheless, selection may induce a correlation between these traits in evolved populations. In Fig. 5a we schematically depict how the raw variation of traits from mutations is distorted by selection and fixation of multiple mutations. Specifically, for a single fixed mutation, selection induces a positive (i.e., antagonistic) correlation between growth rate and lag time. Figure 2a shows this for single fixed mutations, while Fig. 5b, c shows this positive correlation for populations that have accumulated the same number of fixed mutations. We can calculate the Pearson correlation coefficient from the covariation of growth effects *γ* and lag effects *ω* for a single fixed mutation:

**FIG. 5.**
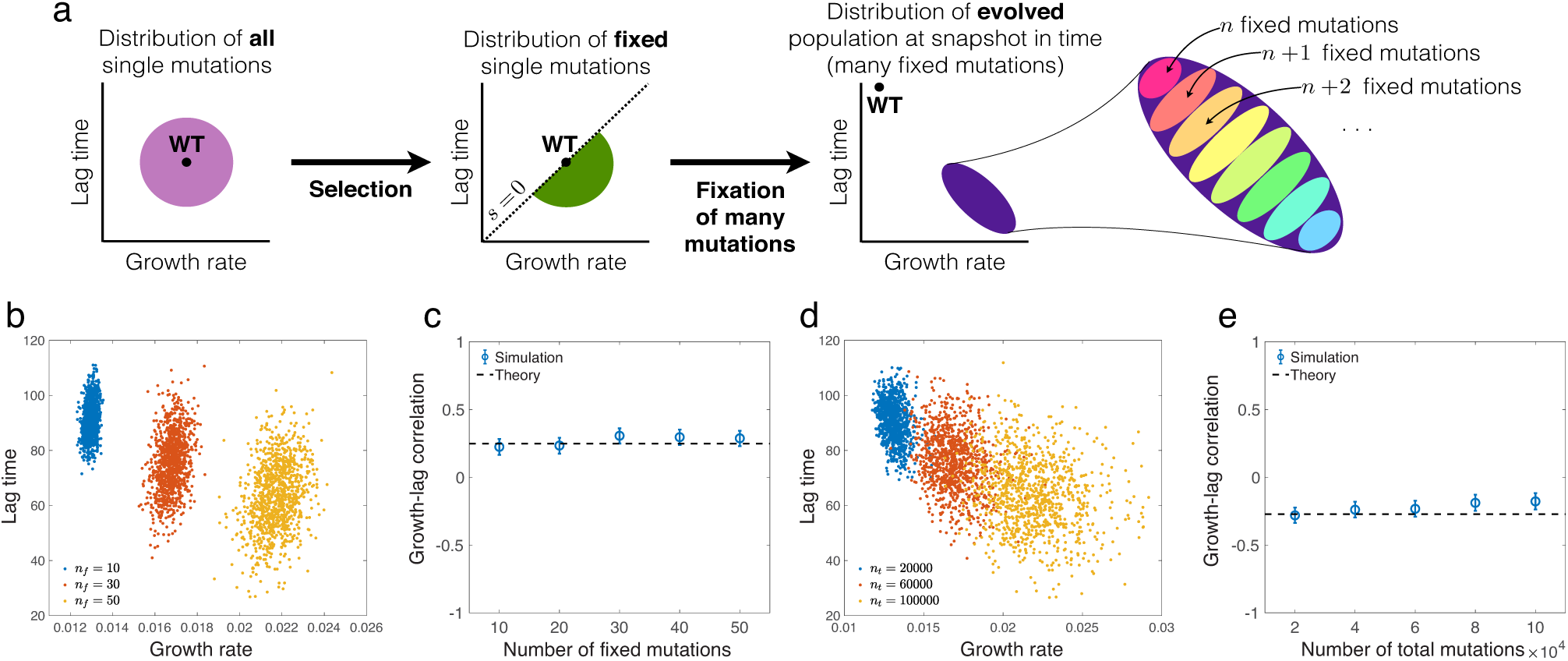
Evolved patterns of covariation among growth traits. (a) Schematic of how selection and fixation of multiple mutations shape the observed distribution of traits. The sign of the Pearson correlation coefficient between the average growth rate and lag time depends on whether we consider an ensemble of populations with the same number of fixed mutations or the same number of total mutation events. (b) Distribution of average growth rate and lag time for 1000 independent populations with the same number of fixed mutations. Each color corresponds to a different number of fixed mutations (*n*_*f*_) indicated in the legend. (c) Pearson correlation coefficient of growth rate and lag time for distributions in panel (b) as a function of the number of fixed mutations. The dashed line is the prediction from Eq. 10 (Supplementary Methods). (d) Same as (b) except each color corresponds to a set of populations at a snapshot in time with the same number of total mutation events. Each color corresponds to a different number of total mutations events (*n*_*t*_) indicated in the legend. (e) Same as (c) but for the set of populations shown in (d). The dashed line is the prediction from Eq. 11 (Supplementary Methods). In (c) and (e) the error-bars represent 95% confidence intervals. In both (b) and (d), we consider the SSWM regime with *D* = 10^3^.

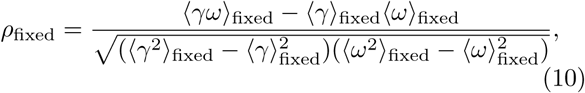

where ⟨·⟩_fixed_ is an average over the distribution of single fixed mutations (Supplementary Methods). We can explicitly calculate this quantity in the SSWM regime, which confirms that it is positive for uncorrelated mutational effects (Supplementary Methods).

However, in evolution experiments we typically observe populations at a particular snapshot in time, such that the populations may have a variable number of fixed mutations but the same number of total mutations that arose (and either fixed or went extinct). Interestingly, the variation in number of fixed mutations at a snapshot in time causes the distribution of growth rates and lag times across populations to stretch into a negative correlation; this is an example of Simpson’s paradox from statistics [43]. Figure 5a shows this effect schematically, while Fig. 5d,e show explicit results from simulations. In Supplementary Methods, we calculate this evolved Pearson correlation coefficient across populations in the SSWM regime to be approximately

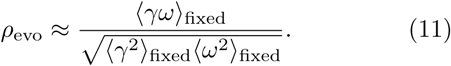

That is, the correlation of traits across populations with multiple mutations is still a function of the distribution of single fixed mutations, but it is not equal to the correlation of single fixed mutations (Eq. 10). Indeed, in Supplementary Methods we explicitly calculate this quantity in the SSWM regime and show that it must be negative for uncorrelated mutational effects.

The predicted correlations in Eqs. 10 and 11 quantitatively match the simulations well in the SSWM regime (Fig. 5c,e). While they are less accurate outside of the SSWM regime, they nevertheless still produce the correct sign of the evolved correlation (Fig. S3a,b,c). However, the signs of the correlations can indeed change depending on the underlying distribution of mutational effects *p*_mut_(*γ, ω, δ*). For example, in Supplementary Methods we explore the effects of varying the mean mutational effects (Fig. S3d) — e.g., whether an average mutation has positive, negative, or zero effect on growth rate — as well as the intrinsic mutational correlation between growth and lag (Fig. S3e).

## DISCUSSION

We have investigated a model of microbial evolution under serial dilution, which is both a common protocol for laboratory evolution experiments [1, 6, 31, 44, 45] as well as a rough model of evolution in natural environments with feast-famine cycles. While there has been extensive work to model population and evolutionary dynamics in these conditions [2, 34, 35, 37], these models have largely neglected the physiological links connecting mutations to selection. However, models that explicitly incorporate these features are necessary to interpret experimental evidence that mutations readily generate variation in multiple cellular traits, and that this variation is important to adaptation [17–20].

In this paper, we have studied a model where mutations can affect three quantitative growth traits — the lag time, exponential growth rate, and yield (Fig. 1a) — since these three traits are widely measured for microbial populations. In particular, we have derived a simple expression (Eq. 3) for the selection coefficient of a mutation in terms of its effects on growth and lag and a single environmental parameter, the dilution factor *D*. While previous work showed that this selection coefficient determines the fixation probability of a single mutation in the SSWM regime [23], here we have shown that this holds even in the presence of clonal interference (Fig. 2b,c,e,f), which appears to be widespread in these experiments [9, 28, 46]. Our result is therefore valuable for interpreting the abundant experimental data on mutant growth traits. We have also calculated the adaptation rates of growth traits per cycle in the SSWM regime, which turn out to increase with the amount of resource *R* and decrease with the dilution factor *D*. These results are confirmed by numerical simulations and remain good predictions even outside of the SSWM regime.

An important difference with the previous work on this model is that here we used a fixed dilution factor *D*, which requires that the bottleneck population size *N*_0_ fluctuates. In contrast, previous work used a fixed *N*_0_ and variable *D* [23, 24]. We observed two important differences between these regimes. First, in the case of fixed *N*_0_ and variable *D*, the fold-change of the population during a single growth cycle 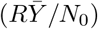 determines the relative selection between growth and lag, since it determines how long the population undergoes exponential growth. Therefore one can experimentally tune this relative selection by varying either the total amount of resources *R* or the fixed bottleneck size *N*_0_. However, when the dilution factor *D* is fixed, the population fold-change is always constrained to exactly equal *D* (Eq. 4; Supplementary Methods), and therefore *D* alone determines the relative selection on growth and lag (Eq. 3). The second difference is that, with fixed *N*_0_ and variable *D*, the selection coefficient depends explicitly on the effective yield 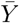 and is therefore frequency-dependent (Supplementary Methods), which enables the possibility of stable coexistence between two strains [23, 24]. However, for the fixed *D* case, the frequency dependence of 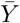 is exactly canceled by *N*_0_ (Eq. 4). Therefore there is only neutral coexistence in this case, requiring the growth and lag traits of the strains to follow an exact constraint set by *D* (Supplementary Methods).

A major result of our model is a prediction on the evolution of covariation between growth traits. In particular, we have shown that correlations between traits can emerge from selection and accumulation of multiple mutations even without an intrinsic correlation between traits from individual mutations (Figs. 5 and S3). We have also shown that selection alone produces no correlation between growth and yield, in the absence of correlated mutational effects (Figs. 2d and 3e). This is important for interpreting evolved patterns of traits in terms of selective or physiological tradeoffs. Specifically, it emphasizes that the evolved covariation between traits conflates both the underlying supply of variation from mutations as well as the action of selection and other aspects of population dynamics (e.g., genetic drift, spatial structure, recombination), and therefore it is difficult to make clear inferences about either aspect purely from the outcome of evolution alone. For example, simply observing a negative correlation between two traits from evolved populations is insufficient to infer whether that correlation is due to a physiological constraint on mutations (e.g., mutations cannot improve both traits simultaneously) or due to a selective constraint (e.g., selection favors specialization in one trait or another).

These questions, of course, have been the foundation of quantitative trait genetics [47]. Historically this field has emphasized polymorphic populations with abundant recombination as are applicable to plant and animal breeding. However, this regime is quite different from microbial populations which, at least under laboratory conditions, are often asexual and dominated by linkage between competing mutations [9, 28, 46]. We therefore need a quantitative description of both between-population as well as within-population covariation of traits of microbial populations in this regime. Recent work has developed some mathematical and simulation results along these lines [48–51], but so far it has not been applied to specific microbial traits.

Microbial growth traits should indeed be an ideal setting for this approach due to abundant data, but conclusions on the nature of trait covariation have remained elusive. Physiological models have predicted a negative correlation between growth rate and lag time across genotypes [52, 53], while models of single-cell variation in lag times also suggests there should be a negative correlation at the whole-population level [54]. However, experimental evidence has been mixed, with some studies finding a negative correlation [13, 16], while others found no correlation [10, 11, 14]. Studies of growth-yield correlations have long been motivated by *r*/*K* selection theory, which suggests there should be tradeoffs between growth rate and yield [55]. For instance, metabolic models make this prediction [56–58]. However, experimental evidence has again been mixed, with some data showing a tradeoff [26, 59, 60], while others show no correlation [15, 18, 19, 61] or even a positive correlation [11, 44]. Some of this ambiguity may have to do with dependence on the environmental conditions [19] or the precise definition of yield. We define yield as the proportionality constant of population size to resource (Eq. 1) and neglect any growth rate dependence on resource concentration. Under these conditions, we predict no direct selection on yield, which means that the only way to generate a correlation of yield with growth rate is if the two traits are constrained at the physiological level, so that mutational effects are correlated. In such cases higher yield could evolve but only as a spandrel [62, 63]. Ultimately, we believe more precise single-cell measurements of these traits, both across large unselected mutant libraries as well as evolved strains, are necessary to definitively test these issues.

## ACKNOWLEDGMENTS

MM was supported by an F32 fellowship from the US National Institutes of Health (GM116217) and an Ambizione grant from the Swiss National Science Foundation (PZ00P3 180147). AA thanks support from NSF CAREER 1752024 and the Harvard Deans Competitive Fund.

