## Supplementary Material for "Evolution of microbial growth traits under serial dilution"

### Supplementary Methods

(Dated: October 16, 2019)

#### I. DETERMINISTIC POPULATION DYNAMICS OVER SERIAL DILUTIONS

At the beginning of the $n$th growth cycle, let the frequencies of each strain $k$ be $x_k(n)$ with total population size $N_0(n)$. To determine the strain frequencies $\{x_k(n+1)\}$ and the initial population size $N_0(n+1)$ for cycle $n+1$, we first note that the selection coefficients relate the frequencies between consecutive cycles according to

$$s_{ij}(n) = \ln\left(\frac{x_i(n+1)}{x_j(n+1)}\right) - \ln\left(\frac{x_i(n)}{x_j(n)}\right), \tag{S1}$$

which follows from the definition in Eq. 2 and the condition that dilution preserves frequencies, i.e., the frequencies at the end of cycle $n$ equal the frequencies at the beginning of cycle $n+1$ (neglecting stochastic effects of sampling). We can rearrange Eq. S1 to determine the frequencies in cycle $n+1$ as functions of the frequencies in cycle $n$ and the selection coefficients:

$$x_i(n+1) = \frac{x_i(n)}{\sum_k x_k(n) e^{s_{ki}(n)}}. \tag{S2}$$

The population size $N_0(n+1)$ for the beginning of cycle $n+1$ is the population size at the end of the $n$th cycle diluted by $D$. The total population size at the end of the $n$th cycle is

$$\begin{aligned}
\sum_k N_0(n) x_k(n) e^{r_k(t_\mathrm{c}(n)-L_k)} &= \left(\frac{R}{N_0(n)\sum_\ell \frac{x_\ell(n)}{Y_\ell} e^{r_\ell(t_\mathrm{c}(n)-L_\ell)}}\right) \sum_k N_0(n) x_k(n) e^{r_k(t_\mathrm{c}(n)-L_k)} \\
&= R\left(\sum_\ell \frac{1}{Y_\ell} \frac{x_\ell(n) e^{r_\ell(t_\mathrm{c}(n)-L_\ell)}}{\sum_k x_k(n) e^{r_k(t_\mathrm{c}(n)-L_k)}}\right)^{-1} \\
&= R\left(\sum_\ell \frac{1}{Y_\ell} \frac{x_\ell(n)}{\sum_k x_k(n) e^{s_{k\ell}(n)}}\right)^{-1} \\
&= R\left(\sum_\ell \frac{x_\ell(n+1)}{Y_\ell}\right)^{-1},
\end{aligned} \tag{S3}$$

where we have inserted the quantity in parentheses on the right-hand side of the first line because it equals 1 according to the saturation equation (Eq. 1), and we invoke Eq. S2 to obtain the last line. Therefore the initial population size in cycle $n+1$ equals this quantity diluted by $D$:

$$N_0(n+1) = \frac{R}{D}\left(\sum_\ell \frac{x_\ell(n+1)}{Y_\ell}\right)^{-1}. \tag{S4}$$

Equation S4 shows that the ratio $R/D$ controls the overall magnitude of the bottleneck population size $N_0(n)$, and hence the effective population size for evolutionary dynamics. Furthermore, Eq. S4 indicates that for $n > 1$, the effective population yield and initial population size are constrained such that

$$\frac{R\bar{Y}(n)}{N_0(n)} = D, \tag{S5}$$

where we define the effective population yield as

$$\bar{Y}(n) = \left(\sum_k \frac{x_k(n)}{Y_k}\right)^{-1}. \tag{S6}$$

#### II. IMPLICIT EQUATION FOR SELECTION COEFFICIENTS

Equation 1 in the main text defines the time $t_\mathrm{c}$ at which the population exhausts the resource and growth stops; Eq. 2 then defines the selection coefficients $s_{ij}$ in terms of $t_\mathrm{c}$. To determine how all $s_{ij}$ depend explicitly on the parameters of the model, we first rewrite Eq. 2 to get $t_\mathrm{c}$ in terms of each $s_{ij}$:

$$t_\mathrm{c} = \frac{s_{ij} + r_i L_i - r_j L_j}{r_i - r_j}. \tag{S7}$$

We then substitute this for $t_\mathrm{c}$ in Eq. 1 and rearrange to obtain an implicit nonlinear equation for the selection coefficients $s_{ij}$:

$$s_{ij} = -\frac{\Delta r_{ij}}{r_i} \ln\left( \frac{N_0}{R} \sum_{\text{strain } k} \frac{x_k}{Y_k} e^{s_{ki}} \right) - \Delta L_{ij} r_j, \tag{S8}$$

where $\Delta r_{ij} = r_i - r_j$ is the difference in growth rates and $\Delta L_{ij} = L_i - L_j$ is the difference in lag times.

#### III. FREQUENCY-DEPENDENT SELECTION AND COEXISTENCE

In general the selection coefficients are frequency-dependent, meaning they depend not only on the traits of the individual strains (lag times $\{L_k\}$, growth rate $\{r_k\}$, and yields $\{Y_k\}$) but also on their frequencies $\{x_k\}$. To find the condition for coexistence of all the strains, we set $s_{ij} = 0$ for all pairs of strains $i$ and $j$ in Eq. S8 and obtain

$$0 = -\frac{\Delta r_{ij}}{r_i} \ln\left( \frac{N_0}{R\bar{Y}} \right) - \Delta L_{ij} r_j, \tag{S9}$$

using the definition for the effective population yield $\bar{Y}$ in Eq. S6. Furthermore, since $R\bar{Y}(n)/N_0(n) = D$ for $n > 1$ (Eq. S5), the dependence on the frequencies $\{x_k\}$ drops out and we obtain

$$\frac{r_i r_j \Delta L_{ij}}{\Delta r_{ij}} = \ln D. \tag{S10}$$

Geometrically, this means that the lag times $\{L_k\}$ and the reciprocal growth rates $\{1/r_k\}$ for all strains must lie on a straight line, with slope $-\ln D$ [1]. If this condition is satisfied by all strains, then the population dynamics are neutral at all frequencies $\{x_k\}$. Conversely, if Eq. S10 is not satisfied, the selection coefficients must be nonzero and because Eq. S10 is independent of the frequencies, the selection coefficient can never change sign. Furthermore, previous work showed that the variation in selection coefficients over the range of frequencies tends to be small [2]. Therefore we can approximate the selection on a strain as its selection coefficient at a low mutant frequency, which we do in the next section.

#### IV. APPROXIMATE SELECTION COEFFICIENT FOR TWO STRAINS

In the two-strain case, we can rewrite the selection coefficient equation (Eq. S8) as

$$s = -\frac{\gamma}{1+\gamma} \ln\left( \frac{N_0}{R} \left[ \frac{1-x}{Y_1} e^{-s} + \frac{x}{Y_2} \right] \right) - \omega, \tag{S11}$$

where $s = s_{21}$ is the selection coefficient of the mutant over the wild-type, $\gamma = (r_2 - r_1)/r_1$ is the relative mutant growth rate, $\omega = (L_2 - L_1) r_1$ is the relative mutant lag time, and $x = x_2$ is the mutant frequency. We approximate the selection coefficient by considering the case of mutant being very rare ($x \to 0$), which is the relevant case for the calculation of the fixation probability [3]. In this case we can exactly solve Eq. S11 to obtain

$$\lim_{x\to 0} s = \gamma \ln\left(\frac{RY_1}{N_0}\right) - \omega(1+\gamma). \tag{S12}$$

We invoke the relation $RY_1/N_0 = D$ over serial dilutions (Eq. S5) and drop higher-order terms in $\gamma$ and $\omega$ to finally obtain (Eq. 3)

$$s \approx \gamma \ln D - \omega. \tag{S13}$$

Alternatively, if we assume the selection coefficient $s$ is small in magnitude, we can expand Eq. S11 in $s$, which yields an identical solution to leading order in $\gamma$ and $\omega$ [1, 2].

#### V. DISTRIBUTION OF MUTATIONAL EFFECTS

When a mutation arises on a background strain with traits $r_1$, $L_1$ and $Y_1$, we randomly generate the new traits $r_2$, $L_2$, and $Y_2$ from a distribution. We assume the mutational effects scale with the values of the background strain's traits, so that the distribution of mutational effects only depends on the relative changes $\gamma = (r_2 - r_1)/r_1$, $\omega = (L_2 - L_1)r_1$, and $\delta = (Y_2 - Y_1)/Y_1$. We ignore epistasis so that mutational effects are additive. In the main text we use a uniform distribution for simplicity:

$$p_{\mathrm{mut}}(\gamma,\omega,\delta) = \begin{cases} (8\gamma_{\max}\omega_{\max}\delta_{\max})^{-1} & \text{for } -\gamma_{\max} < \gamma < \gamma_{\max} \text{ and } -\omega_{\max} < \omega < \omega_{\max} \text{ and } -\delta_{\max} < \delta < \delta_{\max}, \\ 0 & \text{otherwise.} \end{cases} \tag{S14}$$

We use $\gamma_{\max} = 0.02$, $\omega_{\max} = 0.05$, and $\delta_{\max} = 0.02$. In Fig. S3d, we also generalize this uniform distribution by shifting the mean of $\gamma$ and $\omega$ to nonzero values, therefore $-\gamma_{\max} + \mu_\gamma < \gamma < \gamma_{\max} + \mu_\gamma$ and $-\omega_{\max} + \mu_\omega < \omega < \omega_{\max} + \mu_\omega$. We also consider a Gaussian distribution (Fig. S1 and Fig. S3e), with a potentially nonzero Pearson correlation coefficient $\rho_{\mathrm{mut}}$ between growth effects $\gamma$ and lag effects $\omega$:

$$p_{\mathrm{mut}}(\gamma,\omega,\delta) = \frac{1}{(2\pi)^{3/2}\sigma_\gamma\sigma_\omega\sigma_\delta\sqrt{1-\rho_{\mathrm{mut}}^2}} \exp\left(-\frac{1}{2(1-\rho_{\mathrm{mut}}^2)}\left[\frac{\gamma^2}{\sigma_\gamma^2} + \frac{\omega^2}{\sigma_\omega^2} - \frac{2\rho_{\mathrm{mut}}\gamma\omega}{\sigma_\gamma\sigma_\omega}\right] - \frac{\delta^2}{2\sigma_\delta^2}\right). \tag{S15}$$

#### VI. CALCULATION OF THE FIXATION PROBABILITY

To calculate the fixation probability as a function of selection coefficient from the simulations, we discretize the space of relative growth rates $\gamma$ and relative lag times $\omega$ (e.g., Fig. 2a). In each bin we calculate the fixation probability as the ratio between the total number of fixed mutations and the total number of mutations that arose in that bin, across 500 independent populations. We run 5000 cycles in total. To ensure the results are independent of the initial conditions, we collect fixation statistics based on the last 2500 cycles; the results remain the same if we use the last 1250 cycles.

For the uniform distribution of mutational effects (Eq. S14), we use bin sizes of 0.004 for $\gamma$ and 0.01 for $\omega$ (Fig. 2). For the Gaussian distribution (Eq. S15), we use bins of 0.02 for both $\gamma$ and $\omega$ (Fig. S1). Because the ranges of $\gamma$ and $\omega$ of fixed mutations in the Gaussian case are broader than in the uniform case, the resulting fixation probabilities are noisier. However, we do not see a systematic change of the fixation probability as the correlation coefficient between $\gamma$ and $\omega$ varies (Fig. S1). After averaging over the correlation coefficients from $\rho_{\mathrm{mut}} = -0.9$ to $\rho_{\mathrm{mut}} = 0.9$ with a constant interval 0.1 to obtain better statistics, we fit the averaged fixation probability using $\phi_{\mathrm{CI}}(s) = As\exp(-B/s)$ (Eq. 6).

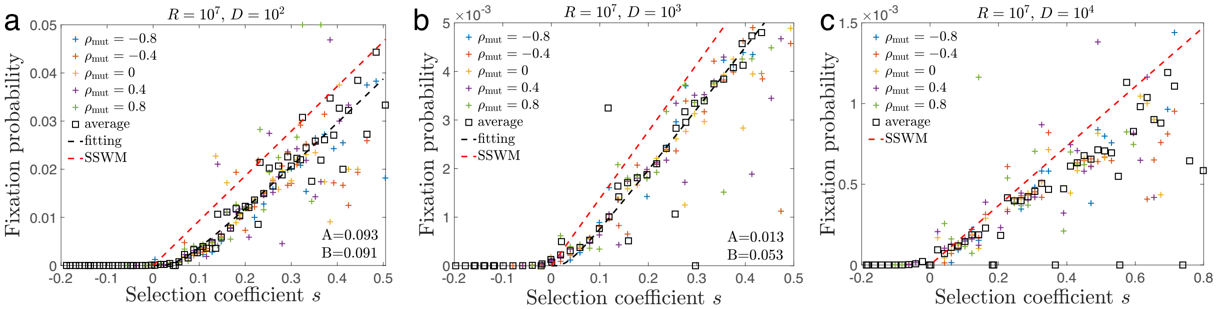


FIG. S1. **Fixation probabilities of Gaussian-distributed mutations.** We test the relation between simulated fixation probability and selection of a mutation relative to its background strain (Eq. 3) for mutational effects that are Gaussian distributed (Eq. S15). Panels (a), (b), and (c) correspond to different values of the resource amount $R$ and dilution factor $D$. Here $\sigma_\gamma = \sigma_\omega = \sigma_\delta = 0.02$. We bin mutations according to the effects $\gamma$ and $\omega$ as described in the text, and for each bin we calculate the simulated fixation probability and the selection coefficient according to Eq. 3. Different colors represent different growth-lag correlation coefficients $\rho_{\text{mut}}$. For (a) and (b), we fit the data to the expression for fixation probability in the clonal interference regime (Eq. 6 in the main text), with the resulting fitting parameters $A$ and $B$ shown in the lower right corner of each panel. The red dashed lines are the prediction of fixation probability in the SSWM regime (Eq. 5 in the main text), which works well for (c).

#### VII. DISTRIBUTION OF FIXED MUTATIONAL EFFECTS IN THE SSWM REGIME

In the SSWM regime, the probability of fixing a mutation with effects $\gamma$ and $\omega$ conditioned on the event of some mutation fixing is

$$P_{\text{fixed}}(\gamma,\omega) = \frac{1}{Z} p_{\text{mut}}(\gamma,\omega)\phi_{\text{SSWM}}(\gamma \ln D - \omega), \tag{S16}$$

where $p_{\text{mut}}(\gamma,\omega)$ is the probability of a mutation with effects $\gamma$ and $\omega$ arising, and the probability of the mutation fixing is (Eq. 5)

$$\phi_{\text{SSWM}}(s) = \frac{2\ln D}{D-1} s\Theta(s), \tag{S17}$$

where $\Theta(s)$ is the Heaviside theta function. We approximate the selection coefficient of the mutation as $s = \gamma \ln D - \omega$ (Eq. 3 or Eq. S13). The normalization factor is $Z$, the probability that a randomly chosen mutation fixes:

$$Z = \int d\gamma \int d\omega \; p_{\text{mut}}(\gamma,\omega)\phi_{\text{SSWM}}(\gamma \ln D - \omega). \tag{S18}$$

To calculate moments of the growth rate effect $\gamma$ and lag time effect $\omega$ of fixed mutations, we must take averages over this distribution. That is, we can calculate the mean value of a function $f(\gamma,\omega)$ as

$$\langle f(\gamma,\omega)\rangle_{\text{fixed}} = \int d\gamma \int d\omega \; P_{\text{fixed}}(\gamma,\omega) f(\gamma,\omega). \tag{S19}$$

##### A. Uniform distribution of mutations

We first consider the case where mutational effects have a uniform distribution (Eq. S14). The normalization factor is

$$Z = \int_{-\gamma_{\max}}^{\gamma_{\max}} d\gamma \int_{-\omega_{\max}}^{\omega_{\max}} d\omega \left(\frac{1}{4\gamma_{\max}\omega_{\max}}\right)\left(\frac{2\ln D}{D-1}\right)(\gamma\ln D - \omega)\Theta(\gamma \ln D - \omega)$$
$$= \begin{cases} \dfrac{3\gamma_{\max}^2 \ln^2 D + \omega_{\max}^2}{6\gamma_{\max}(D-1)} & \text{if } \gamma_{\max}\ln D > \omega_{\max}, \\[2ex] \dfrac{(\gamma_{\max}^2 \ln^2 D + 3\omega_{\max}^2)\ln D}{6\omega_{\max}(D-1)} & \text{if } \gamma_{\max}\ln D < \omega_{\max}. \end{cases} \tag{S20}$$

Therefore the moments of $\gamma$ and $\omega$ are (carrying out integrals in a manner similar to Eq. S20)

$$\langle\gamma\rangle_{\text{fixed}} = \begin{cases} \dfrac{2\gamma_{\max}^3 \ln^2 D}{3\gamma_{\max}^2 \ln^2 D + \omega_{\max}^2} & \text{if } \gamma_{\max}\ln D > \omega_{\max}, \\ \dfrac{2\gamma_{\max}^2 \omega_{\max} \ln D}{\gamma_{\max}^2 \ln^2 D + 3\omega_{\max}^2} & \text{if } \gamma_{\max}\ln D < \omega_{\max}, \end{cases} \tag{S21a}$$

$$\langle\omega\rangle_{\text{fixed}} = \begin{cases} -\dfrac{2\gamma_{\max}\omega_{\max}^2 \ln D}{3\gamma_{\max}^2 \ln^2 D + \omega_{\max}^2} & \text{if } \gamma_{\max}\ln D > \omega_{\max}, \\ -\dfrac{2\omega_{\max}^3}{\gamma_{\max}^2 \ln^2 D + 3\omega_{\max}^2} & \text{if } \gamma_{\max}\ln D < \omega_{\max}. \end{cases} \tag{S21b}$$

$$\langle\gamma^2\rangle_{\text{fixed}} = \begin{cases} \dfrac{15\gamma_{\max}^4 \ln^4 D + \omega_{\max}^4}{10(\ln^2 D)(3\gamma_{\max}^2 \ln^2 D + \omega_{\max}^2)} & \text{if } \gamma_{\max}\ln D > \omega_{\max}, \\ \dfrac{1}{5}\gamma_{\max}^2\left(3 - \dfrac{4\omega_{\max}^2}{\gamma_{\max}^2 \ln^2 D + 3\omega_{\max}^2}\right) & \text{if } \gamma_{\max}\ln D < \omega_{\max}, \end{cases} \tag{S21c}$$

$$\langle\omega^2\rangle_{\text{fixed}} = \begin{cases} \dfrac{1}{15}\omega_{\max}^2\left(5 + \dfrac{4\omega_{\max}^2}{3\gamma_{\max}^2 \ln^2 D + \omega_{\max}^2}\right) & \text{if } \gamma_{\max}\ln D > \omega_{\max}, \\ \dfrac{\gamma_{\max}^4 \ln^4 D + 15\omega_{\max}^4}{10\left(\gamma_{\max}^2 \ln^2 D + 3\omega_{\max}^2\right)} & \text{if } \gamma_{\max}\ln D < \omega_{\max}, \end{cases} \tag{S21d}$$

$$\langle\gamma\omega\rangle_{\text{fixed}} = \begin{cases} -\dfrac{\omega_{\max}^2(5\gamma_{\max}^2 \ln^2 D - \omega_{\max}^2)}{5(\ln D)(3\gamma_{\max}^2 \ln^2 D + \omega_{\max}^2)} & \text{if } \gamma_{\max}\ln D > \omega_{\max}, \\ -\dfrac{1}{5}\gamma_{\max}^2(\ln D)\left(\dfrac{8\omega_{\max}^2}{\gamma_{\max}^2 \ln^2 D + 3\omega_{\max}^2} - 1\right) & \text{if } \gamma_{\max}\ln D < \omega_{\max}. \end{cases} \tag{S21e}$$

We can also calculate the variances and covariances:

$$\langle\gamma^2\rangle_{\text{fixed}} - \langle\gamma\rangle_{\text{fixed}}^2 = \begin{cases} \dfrac{5\gamma_{\max}^6 \ln^6 D + 15\gamma_{\max}^4\omega_{\max}^2 \ln^4 D + 3\gamma_{\max}^2\omega_{\max}^4 \ln^2 D + \omega_{\max}^6}{10(\ln^2 D)\left(3\gamma_{\max}^2 \ln^2 D + \omega_{\max}^2\right)^2} & \text{if } \gamma_{\max}\ln D > \omega_{\max}, \\ \dfrac{3\left(\gamma_{\max}^6 \ln^4 D - 2\gamma_{\max}^4\omega_{\max}^2 \ln^2 D + 5\gamma_{\max}^2\omega_{\max}^4\right)}{5\left(\gamma_{\max}^2 \ln^2 D + 3\omega_{\max}^2\right)^2} & \text{if } \gamma_{\max}\ln D < \omega_{\max}, \end{cases} \tag{S22a}$$

$$\langle\omega^2\rangle_{\text{fixed}} - \langle\omega\rangle_{\text{fixed}}^2 = \begin{cases} \dfrac{3\left(5\gamma_{\max}^4\omega_{\max}^2 \ln^4 D - 2\gamma_{\max}^2\omega_{\max}^4 \ln^2 D + \omega_{\max}^6\right)}{5\left(3\gamma_{\max}^2 \ln^2 D + \omega_{\max}^2\right)^2} & \text{if } \gamma_{\max}\ln D > \omega_{\max}, \\ \dfrac{1}{5}\gamma_{\max}^2(\ln D)\left(1 - \dfrac{8\omega_{\max}^2}{\gamma_{\max}^2 \ln^2 D + 3\omega_{\max}^2}\right) - \dfrac{4\omega_{\max}^6}{\left(\gamma_{\max}^2 \ln^2 D + 3\omega_{\max}^2\right)^2} & \text{if } \gamma_{\max}\ln D < \omega_{\max}, \end{cases} \tag{S22b}$$

$$\langle\gamma\omega\rangle_{\text{fixed}} - \langle\gamma\rangle_{\text{fixed}}\langle\omega\rangle_{\text{fixed}} = \begin{cases} \dfrac{\omega_{\max}^2\left(5\gamma_{\max}^4 \ln^4 D - 2\gamma_{\max}^2\omega_{\max}^2 \ln^2 D + \omega_{\max}^4\right)}{5(\ln D)\left(3\gamma_{\max}^2 \ln^2 D + \omega_{\max}^2\right)^2} & \text{if } \gamma_{\max}\ln D > \omega_{\max}, \\ \dfrac{4\gamma_{\max}^2\omega_{\max}^4 \ln D}{\left(\gamma_{\max}^2 \ln^2 D + 3\omega_{\max}^2\right)^2} + \dfrac{\omega_{\max}^2\left(\omega_{\max}^2 - 5\gamma_{\max}^2 \ln^2 D\right)}{5(\ln D)\left(3\gamma_{\max}^2 \ln^2 D + \omega_{\max}^2\right)} & \text{if } \gamma_{\max}\ln D < \omega_{\max}. \end{cases} \tag{S22c}$$

##### B. Gaussian distribution of mutations

We now repeat the calculation for a Gaussian distribution of mutational effects (Eq. S15). The normalization factor is

$$\begin{aligned} Z &= \int_{-\infty}^{\infty} d\gamma \int_{-\infty}^{\infty} d\omega \left(\frac{1}{2\pi\sigma_\gamma\sigma_\omega}\exp\left(-\frac{\gamma^2}{2\sigma_\gamma^2} - \frac{\omega^2}{2\sigma_\omega^2}\right)\right)\left(\frac{2\ln D}{D-1}\right)(\gamma\ln D - \omega)\Theta(\gamma\ln D - \omega) \\ &= \int_{-\infty}^{\infty} d\gamma \int_{-\infty}^{\gamma\ln D} d\omega \left(\frac{1}{2\pi\sigma_\gamma\sigma_\omega}\exp\left(-\frac{\gamma^2}{2\sigma_\gamma^2} - \frac{\omega^2}{2\sigma_\omega^2}\right)\right)\left(\frac{2\ln D}{D-1}\right)(\gamma\ln D - \omega) \\ &= \frac{2\ln D}{D-1}\sqrt{\frac{\sigma_\gamma^2 \ln^2 D + \sigma_\omega^2}{2\pi}}. \end{aligned} \tag{S23}$$

Therefore the moments of $\gamma$ and $\omega$ are (carrying out integrals in a manner similar to Eq. S23)

$$\langle\gamma\rangle_{\text{fixed}} = \frac{\sigma_\gamma^2 \ln D}{2}\sqrt{\frac{2\pi}{\sigma_\gamma^2 \ln^2 D + \sigma_\omega^2}}, \tag{S24a}$$

$$\langle\omega\rangle_{\text{fixed}} = -\frac{\sigma_\omega^2}{2}\sqrt{\frac{2\pi}{\sigma_\gamma^2 \ln^2 D + \sigma_\omega^2}}, \tag{S24b}$$

$$\langle\gamma^2\rangle_{\text{fixed}} = \sigma_\gamma^2\left(2 - \frac{\sigma_\omega^2}{\sigma_\gamma^2 \ln^2 D + \sigma_\omega^2}\right), \tag{S24c}$$

$$\langle\omega^2\rangle_{\text{fixed}} = \sigma_\omega^2\left(1 + \frac{\sigma_\omega^2}{\sigma_\gamma^2 \ln^2 D + \sigma_\omega^2}\right), \tag{S24d}$$

$$\langle\gamma\omega\rangle_{\text{fixed}} = -\frac{\sigma_\gamma^2\sigma_\omega^2 \ln D}{\sigma_\gamma^2 \ln^2 D + \sigma_\omega^2}. \tag{S24e}$$

The variances and covariances are

$$\langle\gamma^2\rangle_{\text{fixed}} - \langle\gamma\rangle^2_{\text{fixed}} = \sigma^2_\gamma \left(1 - \frac{(\pi-2)\sigma^2_\gamma \ln^2 D)}{2(\sigma^2_\gamma \ln^2 D + \sigma^2_\omega)}\right), \tag{S25a}$$

$$\langle\omega^2\rangle_{\text{fixed}} - \langle\omega\rangle^2_{\text{fixed}} = \sigma^2_\omega \left(1 - \frac{(\pi-2)\sigma^2_\omega}{2(\sigma^2_\gamma \ln^2 D + \sigma^2_\omega)}\right), \tag{S25b}$$

$$\langle\gamma\omega\rangle_{\text{fixed}} - \langle\gamma\rangle_{\text{fixed}}\langle\omega\rangle_{\text{fixed}} = \frac{(\pi-2)\sigma^2_\gamma\sigma^2_\omega \ln D}{2(\sigma^2_\gamma \ln^2 D + \sigma^2_\omega)}. \tag{S25c}$$

#### VIII. ADAPTATION RATES OF THE GROWTH RATE AND LAG TIME IN THE SSWM REGIME

In this section, we calculate the average adaptation speeds of growth rate and lag time using the average changes in these traits determined in Sec. VII. First, the total number of cell divisions in a growth cycle is the population size at the end of the cycle, $N_{\text{final}} = \sum_i N_i(t_c)$, minus the population size at the beginning, $N_0$. We can approximate the final population size as $RY_0$, which assumes that the yields of the evolved strains do not vary significantly from the ancestral yield $Y_0$ (as confirmed by simulations, e.g., Fig. 3e); we also assume $D \gg 1$ so that $N_{\text{final}} - N_0 \approx N_{\text{final}}$. Therefore the total number of mutation events per growth cycle is approximately $\mu RY_0$. For each mutation, the average probability that it fixes is $Z$ (Eq. S18). The expected change in growth rate for a mutation is approximately $\langle\gamma\rangle_{\text{fixed}} r_0$, assuming a small number of fixed mutations so that the growth rate has not changed significantly from the ancestral growth rate $r_0$; similarly, the expected change in lag time is approximately $\langle\omega\rangle_{\text{fixed}}/r_0$.

We find that for both the uniform and Gaussian distributions of mutations, the expected changes in growth rate and lag time per cycle are (Eq. 8)

$$\begin{aligned} W_{\text{growth}} &= \mu RY_0 Z \langle\gamma\rangle_{\text{fixed}} r_0 \\ &= \sigma^2_\gamma r_0 (\ln D) \left(\frac{\mu RY_0 \ln D}{D-1}\right), \end{aligned} \tag{S26}$$

$$\begin{aligned} W_{\text{lag}} &= \mu RY_0 Z \frac{\langle\omega\rangle_{\text{fixed}}}{r_0} \\ &= -\frac{\sigma^2_\omega}{r_0}\left(\frac{\mu RY_0 \ln D}{D-1}\right). \end{aligned} \tag{S27}$$

where $\sigma^2_\gamma = \gamma^2_{\text{max}}/3$ and $\sigma^2_\omega = \omega^2_{\text{max}}/3$ are the variances of $\gamma$ and $\omega$ in the uniform case. The ratio between the growth and lag adaptation speeds defines the average direction of evolution in growth-lag trait space (Eq. 9):

$$\frac{W_{\text{growth}}}{W_{\text{lag}}} = -r_0^2 \frac{\sigma^2_\gamma}{\sigma^2_\omega} \ln D. \tag{S28}$$

We can use this relation to predict the average trajectory of the population growth rate $r_{\text{pop}}$ and lag time $L_{\text{pop}}$ over evolution. In the SSWM regime, we can approximate the average population growth rate and lag time as

$$\langle r_{\text{pop}}\rangle = r_0 + nW_{\text{growth}}, \tag{S29}$$

$$\langle L_{\text{pop}}\rangle = L_0 + nW_{\text{lag}}, \tag{S30}$$

where $n$ is the total number of cycles. Therefore,

$$\frac{\langle r_{\text{pop}}\rangle - r_0}{\langle L_{\text{pop}}\rangle - L_0} = -r_0^2 \frac{\sigma^2_\gamma}{\sigma^2_\omega} \ln D. \tag{S31}$$

In Fig. S2 we compare this equation with the trajectories obtained from simulations for three values of the dilution factor $D$. The prediction matches best for large $D$ (Fig. S2a), since that produces smaller population sizes (through Eq. 4) and therefore better approximates the SSWM limit. The prediction becomes less accurate for small $D$ (Fig. S2b,c) when clonal interference plays a larger role, but still matches well at early times.

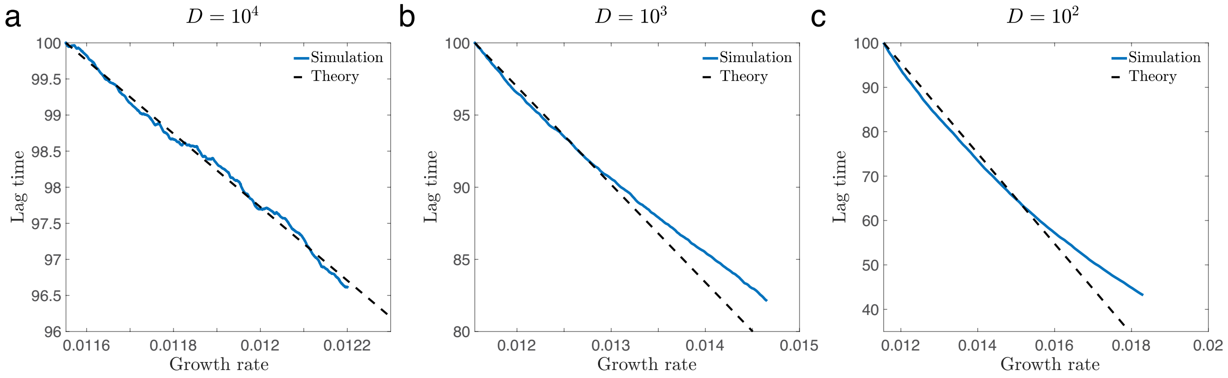


FIG. S2. **Average evolutionary trajectories in growth-lag trait space.** We plot the average population growth rate $r_{\rm pop}$ and lag time $L_{\rm pop}$ from simulations (solid blue lines) with (a) $D = 10^4$, (b) $D = 10^3$, and (c) $D = 10^2$, along with the predicted trajectories in the SSWM limit (Eq. S31; dashed black line).

#### IX. CORRELATION BETWEEN GROWTH RATES AND LAG TIMES

In this section we calculate the evolved correlations between growth rates and lag times. For a single fixed mutation, the correlation coefficient between the relative change in growth rate $\gamma$ and relative change in lag time $\omega$ is

$$\rho_{\rm fixed} = \frac{\langle\gamma\omega\rangle_{\rm fixed} - \langle\gamma\rangle_{\rm fixed}\langle\omega\rangle_{\rm fixed}}{\sqrt{\left(\langle\gamma^2\rangle_{\rm fixed} - \langle\gamma\rangle_{\rm fixed}^2\right)\left(\langle\omega^2\rangle_{\rm fixed} - \langle\omega\rangle_{\rm fixed}^2\right)}}. \tag{S32}$$

However, the quantity more relevant to experimental data is the inter-population correlation between absolute growth rate $r$ and lag time $L$ at a given time when multiple mutations have fixed. To calculate this, we focus on the SSWM regime and assume that each mutation has small effects on the growth rate and lag time, so that the total growth rate $r$ and lag time $L$ can be approximated as sums of these effects:

$$\begin{aligned} r &\approx r_0 + r_0 \sum_{i=1}^{m} \gamma_i, \\ L &\approx L_0 + \frac{1}{r_0}\sum_{i=1}^{m} \omega_i, \end{aligned} \tag{S33}$$

where $r_0$ and $L_0$ are the ancestral growth rate and lag time, and the sums are over all fixed mutations (indexed by $i$) up to the total number $m$.

We can now calculate moments of the evolved growth rate and lag time by averaging over both the distribution of fixed mutations (Eq. S19) and across populations with different numbers of fixed mutations $m$:

$$\begin{aligned} \overline{\langle Lr\rangle_{\rm fixed}} &\approx \frac{1}{M}\sum_{\text{population }\alpha}\left\langle\left(r_0 + r_0\sum_{i=1}^{m_\alpha}\gamma_i\right)\left(L_0 + \frac{1}{r_0}\sum_{i=1}^{m_\alpha}\omega_i\right)\right\rangle_{\rm fixed} \\ &= \frac{1}{M}\sum_{\text{population }\alpha}\left(r_0L_0 + m_\alpha\langle\omega\rangle_{\rm fixed} + L_0r_0m_\alpha\langle\gamma\rangle_{\rm fixed} + m_\alpha\langle\gamma\omega\rangle_{\rm fixed} + (m_\alpha^2 - m_\alpha)\langle\gamma\rangle_{\rm fixed}\langle\omega\rangle_{\rm fixed}\right) \\ &= r_0L_0 + \overline{m}\langle\omega\rangle_{\rm fixed} + L_0r_0\overline{m}\langle\gamma\rangle_{\rm fixed} + \overline{m}\langle\gamma\omega\rangle_{\rm fixed} + (\overline{m^2} - \overline{m})\langle\gamma\rangle_{\rm fixed}\langle\omega\rangle_{\rm fixed}. \end{aligned} \tag{S34}$$

Here the bar indicates an average over all independent populations (total number $M$). Similar calculations yield the (co)variances:

$$\overline{\langle Lr\rangle_{\text{fixed}}} - \overline{\langle L\rangle_{\text{fixed}}}\,\overline{\langle r\rangle_{\text{fixed}}} = \overline{m}\left(\langle \gamma\omega\rangle_{\text{fixed}} - \langle\gamma\rangle_{\text{fixed}}\langle\omega\rangle_{\text{fixed}}\right) + \left(\overline{m^2} - \overline{m}^2\right)\langle\gamma\rangle_{\text{fixed}}\langle\omega\rangle_{\text{fixed}}, \tag{S35}$$

$$\overline{\langle r^2\rangle_{\text{fixed}}} - \left(\overline{\langle r\rangle_{\text{fixed}}}\right)^2 = r_0^2\overline{m}\left(\langle\gamma^2\rangle_{\text{fixed}} - \langle\gamma\rangle_{\text{fixed}}^2\right) + r_0^2\left(\overline{m^2} - \overline{m}^2\right)\langle\gamma\rangle_{\text{fixed}}^2, \tag{S36}$$

$$\overline{\langle L^2\rangle_{\text{fixed}}} - \left(\overline{\langle L\rangle_{\text{fixed}}}\right)^2 = \frac{1}{r_0^2}\overline{m}\left(\langle\omega^2\rangle_{\text{fixed}} - \langle\omega\rangle_{\text{fixed}}^2\right) + \frac{1}{r_0^2}\left(\overline{m^2} - \overline{m}^2\right)\langle\omega\rangle_{\text{fixed}}^2. \tag{S37}$$

That is, the (co)variances of the growth rate and lag time are sums of the (co)variance in the traits for a single fixed mutation and the variance of number of mutations ($\overline{m^2} - \overline{m}^2$). In the SSWM regime, different fixed mutations are independent of each other and the probability of any mutation fixing ($Z$, Eqs. S20 and S23) is small ($Z \sim D^{-1}$ with $D \gg 1$); therefore the number of fixed mutations over a finite time will be approximately Poisson-distributed, so that the variance approximately equals the mean:

$$\overline{m^2} - \overline{m}^2 \approx \overline{m}. \tag{S38}$$

The Pearson correlation coefficient of the evolved growth rate and lag time is therefore (Eq. 11)

$$\begin{aligned}\rho_{\text{evo}} &= \frac{\overline{\langle Lr\rangle_{\text{fixed}}} - \overline{\langle L\rangle_{\text{fixed}}}\,\overline{\langle r\rangle_{\text{fixed}}}}{\sqrt{\left(\overline{\langle r^2\rangle_{\text{fixed}}} - \left(\overline{\langle r\rangle_{\text{fixed}}}\right)^2\right)\left(\overline{\langle L^2\rangle_{\text{fixed}}} - \left(\overline{\langle L\rangle_{\text{fixed}}}\right)^2\right)}} \\ &\approx \frac{\langle\gamma\omega\rangle_{\text{fixed}}}{\sqrt{\langle\gamma^2\rangle_{\text{fixed}}\langle\omega^2\rangle_{\text{fixed}}}}.\end{aligned} \tag{S39}$$

That is, the correlation between evolved growth and lag depends entirely on the moments of growth and lag for a single fixed mutation, but is not identical to the correlation coefficient for a single fixed mutation (Eq. S32).

For the uniform distribution of mutations (Eq. S14), these two correlations equal:

$$\rho_{\text{fixed}} = \begin{cases} \sqrt{\dfrac{2\omega_{\max}^2(5\gamma_{\max}^4\ln^4 D - 2\gamma_{\max}^2\omega_{\max}^2\ln^2 D + \omega_{\max}^4)}{3(5\gamma_{\max}^6\ln^6 D + 15\gamma_{\max}^4\omega_{\max}^2\ln^4 D + 3\gamma_{\max}^2\omega_{\max}^4\ln^2 D + \omega_{\max}^6)}} & \text{if } \gamma_{\max}\ln D > \omega_{\max}, \\ \dfrac{\left(-5\gamma_{\max}^6\omega_{\max}^2\ln^6 D + 31\gamma_{\max}^4\omega_{\max}^4\ln^4 D - 19\gamma_{\max}^2\omega_{\max}^6\ln^2 D + 9\omega_{\max}^8\right)\left(\sqrt{3}\gamma_{\max}(\ln D)\left(3\gamma_{\max}^2\ln^2 D + \omega_{\max}^2\right)\right)^{-1}}{\sqrt{(\gamma_{\max}^4\ln^4 D - 2\gamma_{\max}^2\omega_{\max}^2\ln^2 D + 5\omega_{\max}^4)(\gamma_{\max}^6\ln^5 D - 2\gamma_{\max}^4\omega_{\max}^2\ln^3 D - 15\gamma_{\max}^2\omega_{\max}^4\ln D - 20\omega_{\max}^6)}} & \text{if } \gamma_{\max}\ln D < \omega_{\max}, \end{cases} \tag{S40a}$$

$$\rho_{\text{evo}} = \begin{cases} -\dfrac{\sqrt{2}\omega_{\max}\left(5\gamma_{\max}^2\ln^2 D - \omega_{\max}^2\right)}{\sqrt{\left(5\gamma_{\max}^2\ln^2 D + 3\omega_{\max}^2\right)\left(15\gamma_{\max}^4\ln^4 D + \omega_{\max}^4\right)}} & \text{if } \gamma_{\max}\ln D > \omega_{\max}, \\ -\dfrac{\omega_{\max}^2\left(\gamma_{\max}^2\ln^2 D + 3\omega_{\max}^2\right)\left(5\gamma_{\max}^2\ln^2 D - \omega_{\max}^2\right)}{\gamma_{\max}^2(\ln D)\left(3\gamma_{\max}^2\ln^2 D + \omega_{\max}^2\right)\sqrt{(\ln D)\left(\gamma_{\max}^2\ln^2 D - 5\omega_{\max}^2\right)\left(3\gamma_{\max}^2\ln^2 D + 5\omega_{\max}^2\right)}} & \text{if } \gamma_{\max}\ln D < \omega_{\max}, \end{cases} \tag{S40b}$$

while for the Gaussian distribution of mutations (Eq. S15) they are

$$\rho_{\text{fixed}} = \frac{(\pi - 2)\sigma_\gamma\sigma_\omega\ln D}{\sqrt{\left[(4-\pi)\sigma_\gamma^2\ln^2 D + 2\sigma_\omega^2\right]\left[2\sigma_\gamma^2\ln^2 D + (4-\pi)\sigma_\omega^2\right]}}, \tag{S41a}$$

$$\rho_{\text{evo}} = -\frac{\sigma_\gamma\sigma_\omega\ln D}{\sqrt{2\sigma_\gamma^4\ln^4 D + 5\sigma_\gamma^2\sigma_\omega^2\ln^2 D + 2\sigma_\omega^4}}. \tag{S41b}$$

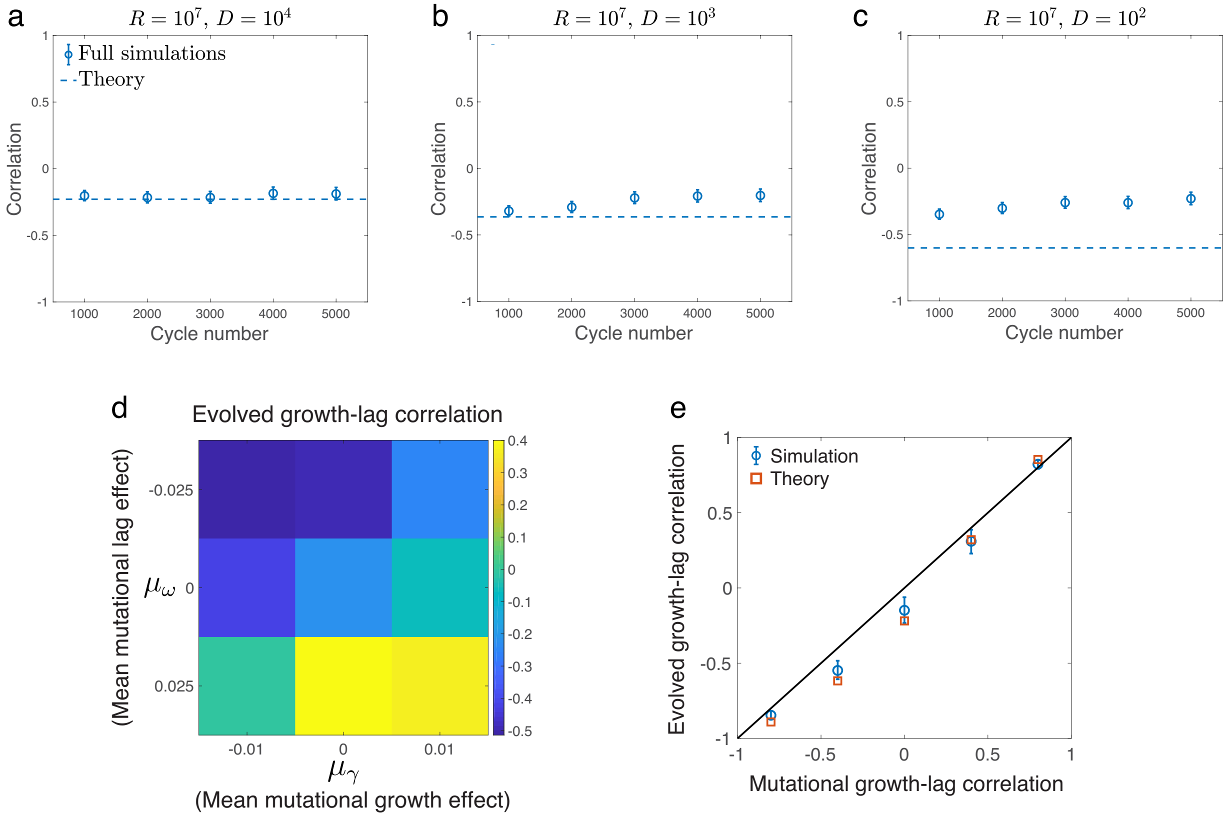


FIG. S3. **Evolved patterns of covaration among growth traits.** (a-c) Pearson correlation coefficients of the population-averaged growth rates and lag times versus the cycle number from simulations. The blue circles are the measured values from the full simulations and the dashed lines are the predictions for the SSWM regime (Eq. 11). The error bars represent 95% confidence intervals. (d) Evolved correlation coefficient $\rho_{\mathrm{evo}}$ of growth rate and lag time (after 50000 mutational trials) as a function of the mean mutational effects on growth rate and lag time (Eq. S14). (e) Evolved correlation coefficient $\rho_{\mathrm{evo}}$ of growth rate and lag time (after 50000 mutational trials) as a function of the mutational correlation $\rho_{\mathrm{mut}}$ of these two traits (Eq. S15). The blue points show simulation results, while the red points show the prediction from Eq. 11. The black line shows the line of identity. In both (d) and (e), we consider the SSWM regime with $D = 10^3$.

Note that the correlation $\rho_{\mathrm{fixed}}$ for a single fixed mutation is always positive, while the correlation $\rho_{\mathrm{evo}}$ between evolved traits is always negative (cf. Fig. 5).

[1] M. Manhart and E. I. Shakhnovich, Nat Commun **9**, 3214 (2018).
[2] M. Manhart, B. V. Adkar, and E. I. Shakhnovich, Proc R Soc B **285** (2018).
[3] Y. Guo, M. Vucelja, and A. Amir, Science Advances **5** (2019), 10.1126/sciadv.aav3842.
